# A Systems-Level Transcriptomic Analysis of Polycystic Ovary Syndrome as a Mitochondrial-Immunometabolic Disorder

**DOI:** 10.64898/2026.01.22.701018

**Authors:** Ritika Patial, Sonalika Ray, Kashmir Singh, R.C. Sobti

## Abstract

Polycystic Ovary Syndrome (PCOS) is known as an endocrine and metabolic disorder; however, emerging molecular evidence suggests a far more complex systems-level pathology. In this study, we performed an integrative transcriptomic and pathway-level analysis of endometrial tissue from women with PCOS to gain a deeper understanding of the underlying mechanism facilitating the disorder. The findings of the study highlighted mitochondrial dysfunction, chronic oxidative stress, and multi-layered immune dysregulation, adding some new insight apart from classical hyperandrogenism and insulin resistance. We identified some novel gene disease associations which involve ***C15orf48, ODF3B PRR15-DT, LINC01176,*** and ***LOC105379193***. The upstream regulators such as (**NFE2L2, TWNK, ALKBH1, BCOR, SMARCA4**) involved in processes including mitochondrial genome, redox balance, and chromatin remodeling provided new insights into regulatory mechanisms. The IPA pathway analysis validated the compromised immune recovery with low grade inflammations and mitochondrial dysfunctionality. The observations emphasize on complex associations discarding its PCOS pure endocrine nature through immunometabolic-mitochondrial dysfunctionalities.

## 1. Introduction

With the significant shift in lifestyle of the individuals, Polycystic Ovary Syndrome (PCOS), a heterogeneous condition, has become highly prevalent and affects 6–15% of women of reproductive age (1,2). The classic symptoms include hyperandrogenism, ovulatory dysfunction, and a close association with insulin resistance, obesity, and metabolic syndrome (3). Apart from its significant role in reproductive dysfunction, PCOS is commonly associated with long-term comorbidities, including type 2 diabetes mellitus, cardiovascular disease, endometrial hyperplasia, and an increased risk of endometrial cancer (4). The higher rates of infertility, implantation failure, and recurrent pregnancy loss are also experienced by women with PCOS, highlighting the broader impact of PCOS on women’s physical, social, and mental well-being (5). The heterogeneity of PCOS as according to Rotterdam diagnostic criteria makes diagnosis and management more complicated (6). Therefore, it emphasizes the need for studies that explore tissue-specific and phenotype-specific differences.

Despite extensive research, the pathophysiology of PCOS remains unexplored, and clinical management continues to rely largely on symptomatic treatment rather than targeted. Several studies also highlight the fact that PCOS involves more than endocrine abnormalities, such as metabolic stress, mitochondrial dysfunction, and immune dysregulation. The role of mitochondria is not only restricted to energy metabolism but has also been seen in regulating inflammatory signaling and oxidative stress responses. The activation of inflammatory pathways through excessive production of reactive oxygen species (ROS) is also a known contributing factor to chronic low-grade inflammation observed in PCOS across multiple tissues (7). Numerous findings involving mitochondrial dysfunction, elevated reactive oxygen species (ROS), chronic low-grade inflammation, and dysregulated immune signaling across ovarian, adipose, and endometrial tissues have been made (8,9). These findings hint towards complex interactions between metabolic stress, immune dysregulation, and metabolic imbalance. The persistent presence of inflammatory and oxidative stress may drive pathological tissue remodeling, particularly through excessive extracellular matrix (ECM) deposition and fibrosis. Such remodeling highlights a continuous injury-repair response that affects tissue architecture and its communication. Additionally, structural remodeling through extracellular matrix (ECM) accumulation and fibrosis endometrial stroma has also been reported(10). This modelling points towards a constant tissue-level response to chronic inflammation and oxidative stress. Most studies on PCOS have focused on ovarian tissue or blood, but very few have looked closely at the endometrium (11). The endometrium is a dynamic tissue strongly affected by the coordinated working of metabolic, hormonal, and immune signals to support embryo implantation and pregnancy. (12). Disruption of this coordination can impact the receptivity of endometrium, leading to implantation failure and also some unfavorable reproductive conditions. Several reports into dysregulation of pathways related to oxidative phosphorylation, chemokine signaling, lipid metabolism, and steroid biosynthesis have been made through transcriptomics studies. Despite advancing our understanding of PCOS, yet the majority of investigations only focus on individual pathways in isolation or on protein-coding genes and we still do not have a clear understanding of how mitochondrial changes, immune activity, and tissue remodeling work together in the endometrium. To fill these gaps, our study uses integrated gene expression and pathway analyses on endometrial tissue from women with PCOS. By identifying these transcriptomic changes, we aim to provide a more complete, systems-level understanding of PCOS.

## 3. Research Methodology

### 2.1 Data Extraction

The raw gene expression data for this study was obtained from the NCBI-GEO GSE277906, which included a total of 40 samples, with 23 from women diagnosed with PCOS and 17 from healthy controls (13). The raw obtained data was preprocessed, and duplicate gene symbols, genes with zero or very low expression values, and genes with no measurable variance across samples were removed to obtain significant genes with measurable differences in expression between PCOS and normal samples. Following the pre-processing step, the next step was to transform the expression matrix using log transformation to stabilize variance and to prevent any outliers.

### 2.3 Exploratory Data Analysis

The log-transformed data was used for principal component analysis to assess the variability in the dataset and to look for any potential outliers. PCA is a dimensionality reduction technique that reduces the dimensionality of the data to make visualization of the variation of points in the data easier (14).

### 2.4 Differential Gene Expression Analysis

DGE analysis between PCOS and normal samples using the DESeq2 package was performed on raw data(15). To highlight the most statistically significant DEGs, strict filtering criteria was applied of p-value < 0.05 and Benjamini–Hochberg false discovery rate correction was applied for multiple testing < 0.05 (16). Further we used Log2 fold change of ≥ 2 or <-2 to focus on high-effect genes. Further, the volcano plot was created using the EnhancedVolcano package in R. Genes absent from CTD PCOS-curated lists and lacking PCOS-related publications in PubMed (search: ‘gene name AND PCOS’ up to June 2025) were labeled as putative novel associations.

### 2.3 Functional Characterization

To understand the biological relevance of the DEGs, functional enrichment analysis using the ClusterProfiler package in R was performed (17). Also, to understand the associated biological processes (BP), molecular functions (MF), and cellular components (CC) associated with the DEGs (18), Gene Ontology (GO) enrichment was performed. In addition, KEGG pathway enrichment analysis was performed to understand the involvement of DEGs in known signaling and metabolic pathways (19).

### 2.4 Ingenuity Pathway Analysis (IPA)

IPA (QIAGEN, version 2024.2; knowledge base updated March 2024) was used with default setting for analysis on the DEGs to get deeper insights through Canonical Pathways, Diseases and Biofunctions, and Upstream Regulators (20). Canonical pathways and Diseases and Biofunctions using a threshold of < 0.05 for p-value, z-scores of ≥ 2 and <-2 for activated and inhibited pathways were obtained. In the upstream regulator analysis, transcription factors, cytokines, and signaling molecules potentially responsible for the observed gene expression changes were identified with the filtering criteria of z-scores ≥ 2 or <-2 and a significant p-value.

### 2.7 Systems Biology Approach and Network Construction

To gain deeper insight into the molecular interactions among the DEGs, a systems biology approach using protein-protein interaction (PPI) network analysis was done using STRING database and then visualization was done in Cytoscape (21). To focus only on the significant interactions, a confidence score of 0.7 was used.

### 2.8 GSEA analysis

Gene Set Enrichment Analysis (GSEA) was performed to identify significantly enriched biological pathways associated with the DEG using the **fgsea** package in multilevel mode, with gene sets obtained from **MSigDB via msigdbr** for *Homo sapiens*.(23). Two MSigDB collections were analyzed: **Hallmark (H)** and **Reactome pathways (C2:CP:REACTOME)**. Genes were ranked using a composite metric which includes the direction and magnitude of differential expression together with statistical significance. Adjusted *p*-values were used when available; otherwise, nominal *p*-values were applied. When present, differential expression test statistics (e.g., Wald or *t* statistics) were used directly for gene ranking. Only genes overlapping with the selected gene sets were included. Enrichment significance was evaluated using fgsea’s **adaptive multilevel algorithm**, which does not rely on a fixed number of permutations. Pathways were considered significantly at a **false discovery rate (FDR) < 0.25**, with an additional filter of **FDR < 0.05**.

## Results

### Principal Component Analysis

**Figure 1 A:**
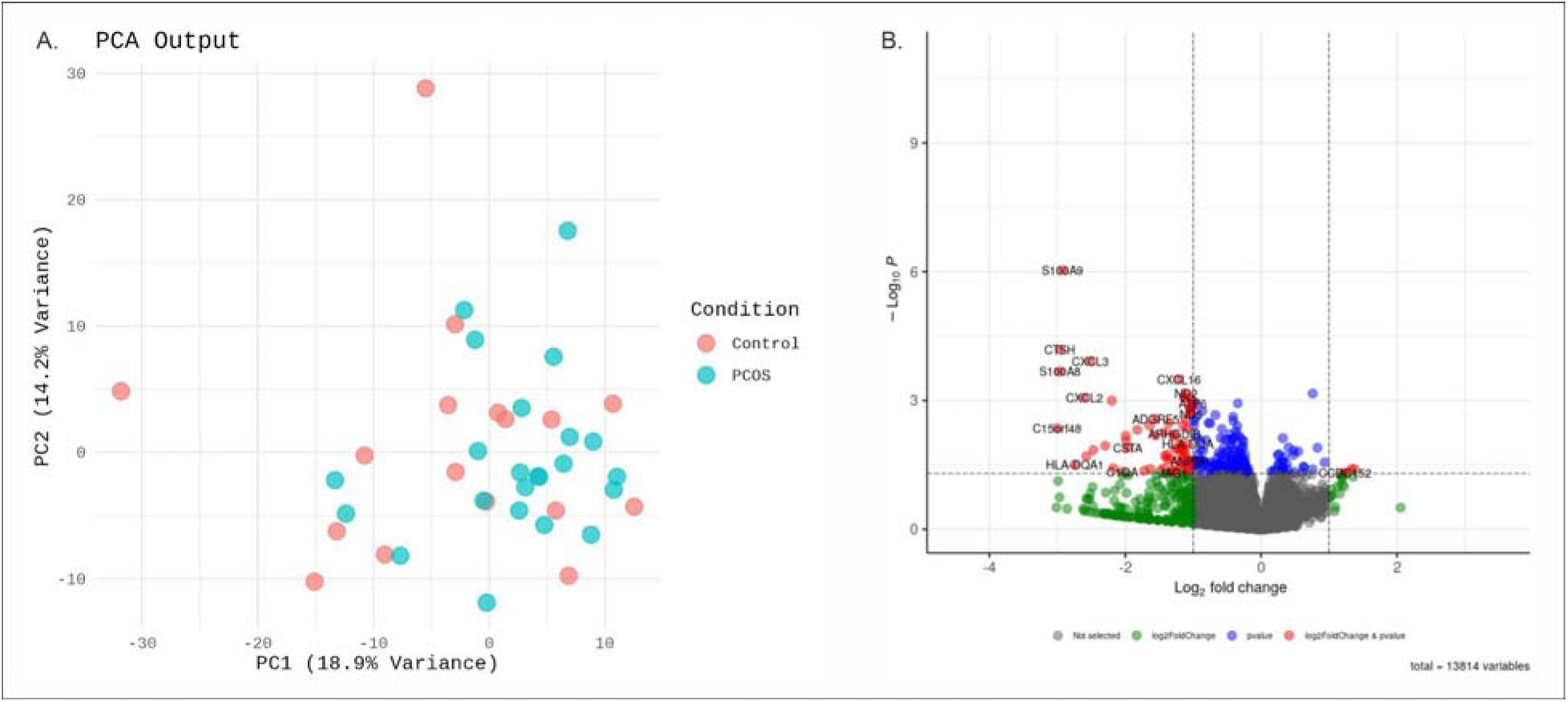
The plot represents the 2-dimensional PCA plot of the PCOS samples, showing the variance among them. The blue ring represents the cluster of PCOS samples, and the red ring represents the control cluster.

The PCA plot shown in figure 1A represents the variance in gene expression between PCOS and control (CN) samples, with PC1 accounting for 23.3% of the variance and PC2 contributing 10.9%, together explaining 34.2% of the total variance in the dataset. It shows some separation between PCOS and CN groups and the partial overlap indicates shared expression patterns, suggesting that while PCOS has distinct molecular characteristics, some genes exhibit similar expression in both conditions.

#### Differentially Expressed Gene Analysis

From differential gene expression analysis, a total of 75 significant DEGs were identified at a p.adj < 0.05 and absolute log□ fold change (|log□FC|) > 2, figure 1B. Out of the total, 4 were upregulated, and 71 were downregulated genes. Among the top significantly upregulated genes were *COX2, COX3, DEF6*, and *ND1* and downregulated genes included *S100A9, CTSH, CXCL3, S100A8, CXCL16* amongst others. Further, in comparison with the gene list from Comparative Toxicogenomics Database (CTD) we identified 12 DEGs that did not have known, curated therapeutic or marker/mechanism roles directly associated with PCOS **supplementary table 1**. Those genes included *ADGRE5, ARHGAP45, BOLA2B, LINC01176, LOC105379193, ND4L, P3H3, PRR15-DT, SQOR, TAMALIN, C15orf48,* and *ODF3B*. Further filtering through literature with PubMed searches using gene names combined with disease-specific terms for polycystic ovary syndrome, including synonymous terminology. Searches incorporated Title/Abstract fields and MeSH terms and were filtered to include only human studies in female populations, we identified 5 genes namely *C15orf48,* and *ODF3B PRR15-DT, LINC01176, LOC105379193* as novel associations with PCOS. two mitochondrial genes namely ND4L (NADH dehydrogenase subunit 4L) and SQOR (sulfide quinone oxidoreductase) essential for oxidative phosphorylation and redox equilibrium (24)(25). Their significant downregulation in the PCOS samples highlights an impairment in mitochondrial bioenergetics and sulfur metabolism, which are critical factors in the oxidative stress and insulin resistance. Likewise, ADGRE5 (CD97) (Adhesion G Protein-Coupled Receptor E5) (26) and ARHGAP45 (Rho GTPase-activating protein 45) (27) showed reduced expression, indicating a dysregulation in immune cell adhesion and cytoskeletal remodeling processes (28). The significant downregulation of C15orf48 logfc −2.19877, a modulator of Cytochrome C Oxidase during Inflammation (29), is a gene associated with the control of inflammatory responses. Coding and non-coding roles of MOCCI (C15ORF48) coordinate to regulate host inflammation(30) (31). Alongside these protein-coding genes, other non-coding RNA transcripts demonstrated notable differential expression (32). The long non-coding RNAs (lncRNAs) such as LINC01176, PRR15-DT, and LOC105379193 known for their regulatory functions in chromatin organization and transcriptional regulation. Their altered expression levels suggest possible new insight of epigenetic control of ovarian function and metabolic signaling networks in POCS. BOLA2B (Iron-Sulfur Cluster Formation and Cellular Transport) (33), ODF3B (Cellular Architecture and Proliferation) (34), and TAMALIN (Post-Synaptic Scaffolding and Cytoskeletal Dynamics) not known to be associated with reproductive or metabolic issues, but their presence as differentially expressed genes suggests that they may be involved with cellular transport systems, mitochondrial architecture, and tissue remodeling processes (35) (36). The downregulation of P3H3 (Prolyl 3-Hydroxylase 3), an enzyme critical for collagen hydroxylation and maintenance of extracellular matrix, suggest a possible changes in stromal structure and stromal-epithelial signaling that could impact endometrial receptivity and normal tissue homeostasis (37).

#### Gene Ontology (GO) Enrichment Analysis

To understand the biological significance of the DEGs, a Gene Ontology (GO) enrichment analysis was performed focusing on Biological Process (BP), Molecular Function (MF), and Cellular Component (CC). GO Biological Process (BP) figure 2A and 2B.

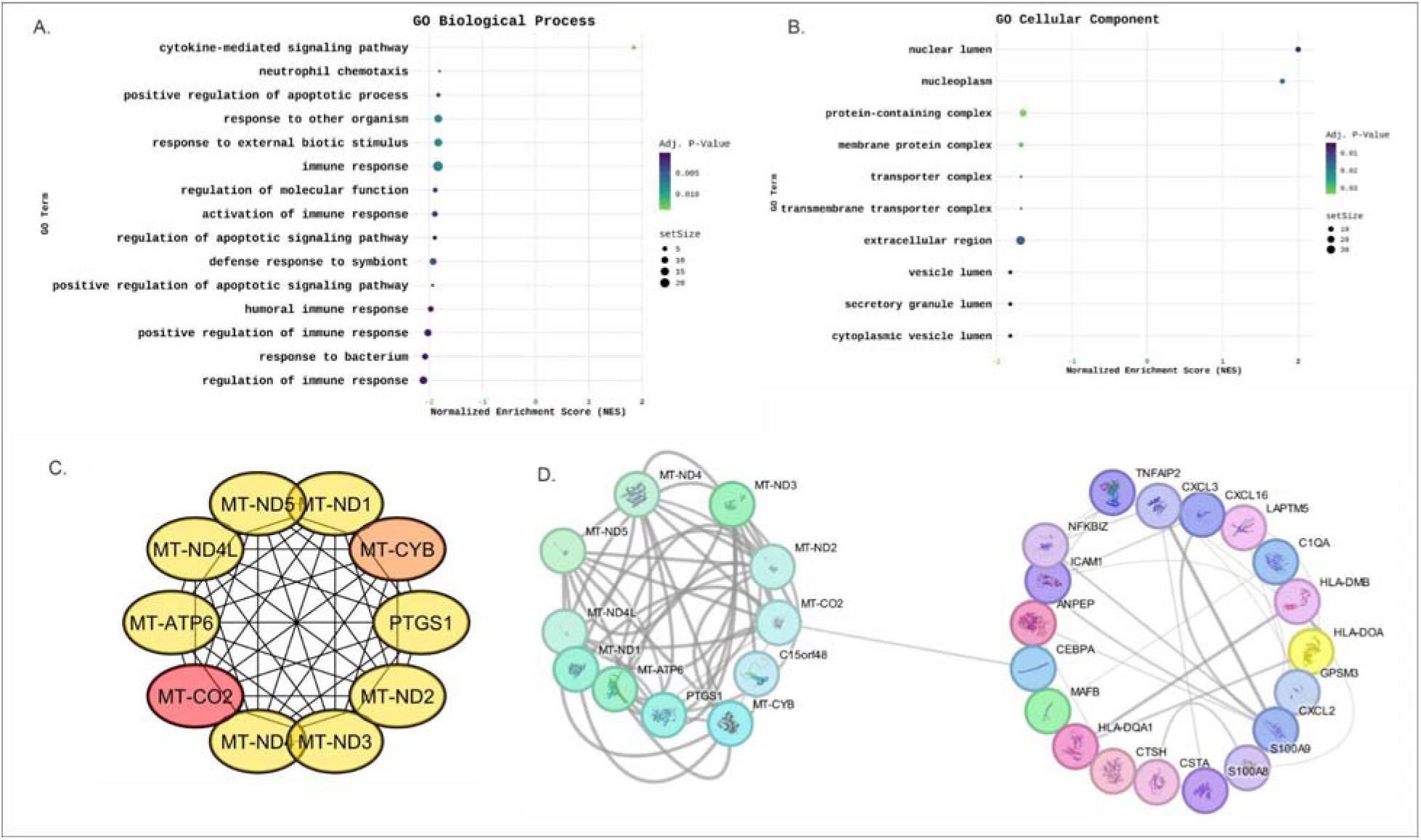

The GO-BP includes significant biological processes and signaling pathways associated with disease pathophysiology. The analysis highlights enrichment of immune-related and cytokine-mediated pathways, highlighting the immune-modulatory landscape of PCOS. Key pathways include lymphocyte and T cell differentiation (33), cytokine-mediated signaling, and responses to peptide or cytokine stimuli (34), emphasizing immune modulation and inflammatory regulation, indicating active immunological adaptation and transcriptional reprogramming within the tissue environment. Furthermore, the processes related to molecular mediator manufacturing and post-translational protein modification were also enriched highlighting the intricate regulation of immunological signaling and protein functionality. Biological processes collectively state that PCOS endometrial tissue experiences immune activation, cytokine-mediated signaling, and also differentiation modification.

##### GO Cellular Component (CC)

The GO-CC highlights the DEGs to be situated within nuclear and membrane-associated compartments, surrounding protein-containing, membrane protein, and transmembrane transporter complexes, along with the nucleoplasm, and the nuclear lumen. The enrichment of nuclear domains, which includes nucleoplasm and nuclear lumen, signifies the role of these genes in transcriptional control, chromatin architecture, and gene regulatory mechanisms, suggesting substantial nuclear remodeling and transcriptional reprogramming in PCOS.

##### GO Molecular Function (MF)

The Gene Ontology (GO) Molecular Function (MF) includes immunological and metabolic signaling pathways, such as calcium-dependent protein binding, immune receptor activity, RAGE receptor binding (27), zinc ion binding, and lipid binding. These molecular processes are known to regulate signal transduction, inflammation, and oxidative stress detection. The activation of RAGE receptor binding signifies the activation of redox and inflammatory signaling. This also aligns with oxidative stress-induced immunological regulation in PCOS. Also, Zinc and lipid binding similarly suggest that there are disturbances in metal-ion equilibrium and lipid metabolism., crucial to insulin resistance and chronic inflammation in the pathogenesis of PCOS.

#### IPA Analysis

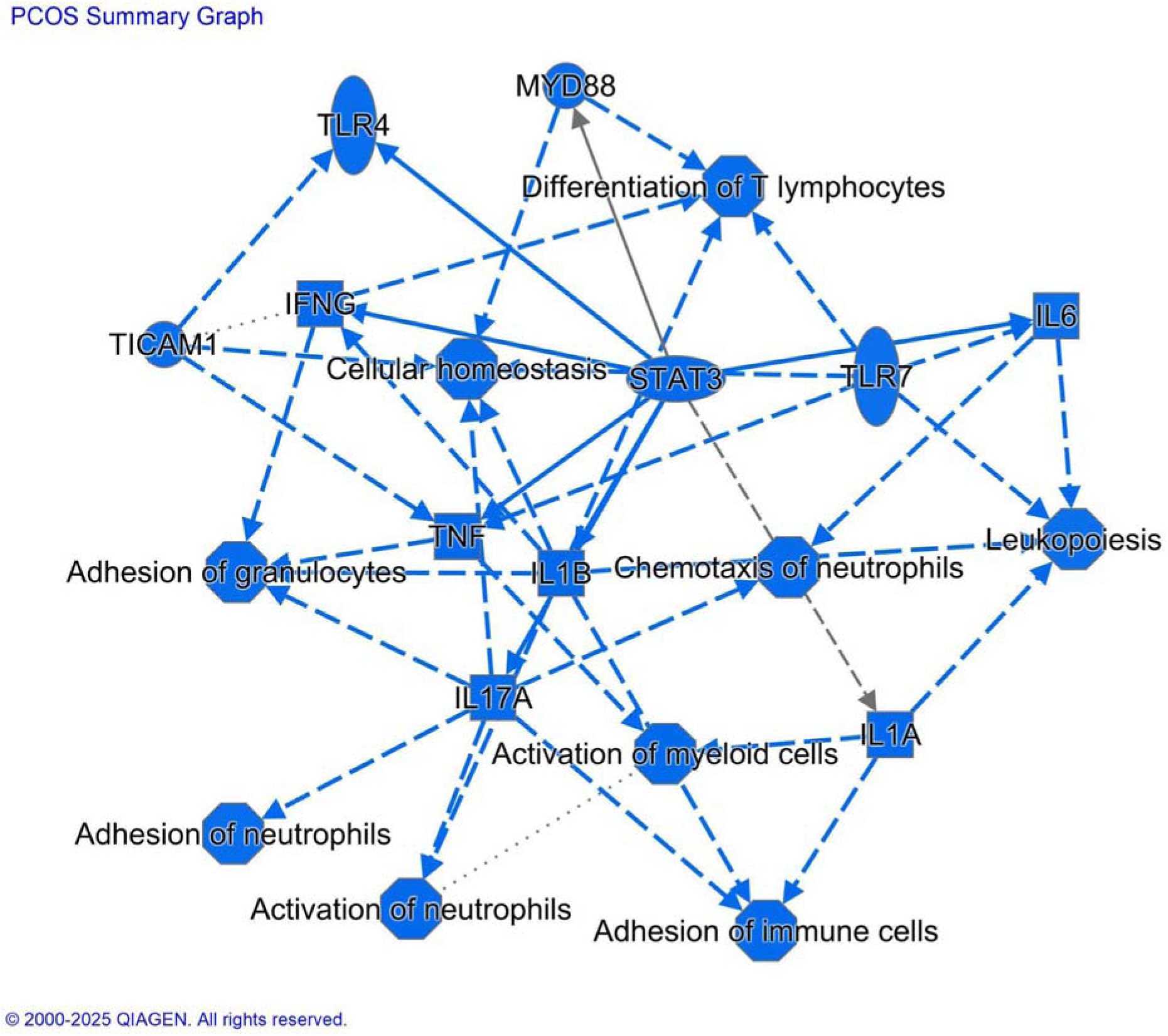

The graphical summary emphasizes the highly **interconnected inflammatory and immune signaling network** in the PCOS endometrium. The network contributes to altered immune interactions and tissue homeostasis affecting endometrial receptivity. **STAT3** appears as a central regulatory node, integrating signals from multiple upstream inflammatory mediators, including **IL6, IL1B, TNF, IFNG**, and **IL17A**, indicating the activation of cytokine-driven signaling pathways. Upstream regulators such as **TLR4** and **TLR7**, and **MYD88**, suggest activation of **innate immune sensing pathways**. The activation further contributes to downstream cytokine production and immune cell recruitment. The network highlights several biological processes related to **leukopoiesis**, **chemotaxis of neutrophils**, **activation and adhesion of myeloid and immune cells**, and **adhesion of granulocytes**, pointing to an inflammatory microenvironment dominated by innate immune responses. The **IL17A-IL1B-TNF** signaling nodes, together with neutrophil-related functions, support a **pro-inflammatory, neutrophil-driven immune axis**. **IFNG** and **T-lymphocyte differentiation** indicate involvement of adaptive immune components. These immune processes converge on **cellular homeostasis**, suggesting that sustained immune activation may directly impact the cellular stability of endometrial tissue along with its function.

#### Upstream regulators

Upstream regulator analysis highlighted several prominent regulators such as RSAD2, NQO1, SIRT3, STAT3, SREBF1, FBXW7, EGF, DNMT3A, LEP, JUN, TGFBR2, STAT1, and AGO2, known to be previously associated with polycystic ovarian syndrome (PCOS). These regulators collectively represent the pivotal role of oxidative stress, lipid metabolism, and cytokine signaling in the pathogenesis of PCOS. SIRT3 and NQO1 are part of mitochondrial redox control and antioxidant defense (24) (25), whereas STAT3 and STAT1 regulate inflammatory transcriptional responses and cytokine feedback mechanisms (26). The presence of SREBF1 and LEP points to the metabolic aspect of PCOS, connecting lipid synthesis and adipokine signaling to ovarian and systemic dysfunction. In addition to these established regulators, several additional upstream effectors have been identified that are not associated with the molecular pathophysiology of PCOS **table 1**. It includes ALKBH1, TWNK, BCOR, TFE3, HIBCH, PPARGC1B, NFE2L2, KLF6, NFKB2, and SMARCA4. The functional annotation of these regulators suggests their role in mitochondrial genome preservation, chromatin restructuring, and transcriptional stress responses. TWNK (Twinkle helicase), crucial for mitochondrial DNA replication (32), indicates disruption in mitochondrial genome integrity and bioenergetic equilibrium, along with the established mitochondrial malfunction in granulosa and theca cells of PCOS. ALKBH, a dioxygenase involved in nucleic acid demethylation, connecting metabolic intermediates to epigenetic regulation (33), is a possible mechanism for the metabolic-epigenetic interaction typical of PCOS. BCOR, a transcriptional corepressor, regulates developmental signaling and chromatin remodeling, and its activation suggests inhibition of mitochondrial transcription and modified cell differentiation signals identified in the samples.

PPARGC1B (PGC-1β), a transcriptional coactivator of nuclear receptors known in oxidative phosphorylation (44), marks mitochondrial biogenesis and metabolic adaptability as key regulatory targets. SMARCA4 (BRG1), a chromatin-remodeling ATPase, and KLF6 (Krüppel-like factor 6), are the transcription factors responsive to stress, indicating extensive nuclear reprogramming associated with metabolic stress and inflammation. HIBCH, an enzyme involved in valine catabolism, indicates secondary metabolic disruptions associated with amino acid metabolism and mitochondrial redox equilibrium. NFKB2 was recognized as an additional regulator known to regulate inflammatory and immunological responses, linking mitochondrial failure with persistent inflammation, a characteristic of PCOS (26).

NFE2L2 has emerged as a crucial transcriptional regulator among the new regulators. NFE2L2 regulates antioxidant and cytoprotective gene networks in response to oxidative stress (45). The analysis highlighted that NFE2L2 regulates C15orf48, a mitochondrial-associated protein-coding gene with an unclear function. The activation of C15orf48 and NFE2L2 indicates a new regulatory axis controlling mitochondrial stress adaptation and immunological modulation. C15orf48 is reported as an interferon-inducible factor situated in mitochondria, regulating innate immune signaling and apoptosis (30) (46). The coordinated alteration of **NFE2L2 activity and C15orf48** suggests the existence of a regulatory axis. The axis may link **impaired antioxidant defense with mitochondrial stress signaling and immune modulation**, potentially contributing to an oxidative stress-prone and inflammatory endometrial microenvironment. A kinase receptor TGFBR2 (transforming growth factor beta receptor II), known to be associated with fibrosis and ovarian follicular development, was discovered to be linked with C15orf48. The interaction between NFE2L2 and TGFBR2 signaling reveals an unknown association between oxidative stress regulation and TGF-β-mediated fibrotic remodeling, which may reveal granulosa cell failure and disrupted folliculogenesis in PCOS figure 3. The targeting of C15orf48 by (NFE2L2) and a kinase (TGFBR2) highlights linking oxidative homeostasis, immunological control, and reproductive tissue remodeling. Canonical regulators such as SIRT3, STAT3, and TGFBR2 support the association of oxidative stress and inflammatory signaling to PCOS. On the other hand, the discovery of novel upstream molecules NFE2L2, BCOR, TWNK, and ALKBH1 provides new molecular insights into mitochondrial genome regulation, redox homeostasis, an chromatin-mediated transcriptional adaptation. The identification of the NFE2L2-C15orf48-TGFBR2 axis highlights an overlooked molecular connection between antioxidant response and reproductive tissue remodeling, underscoring a potential pathway for future therapeutic investigation in PCOS (45).

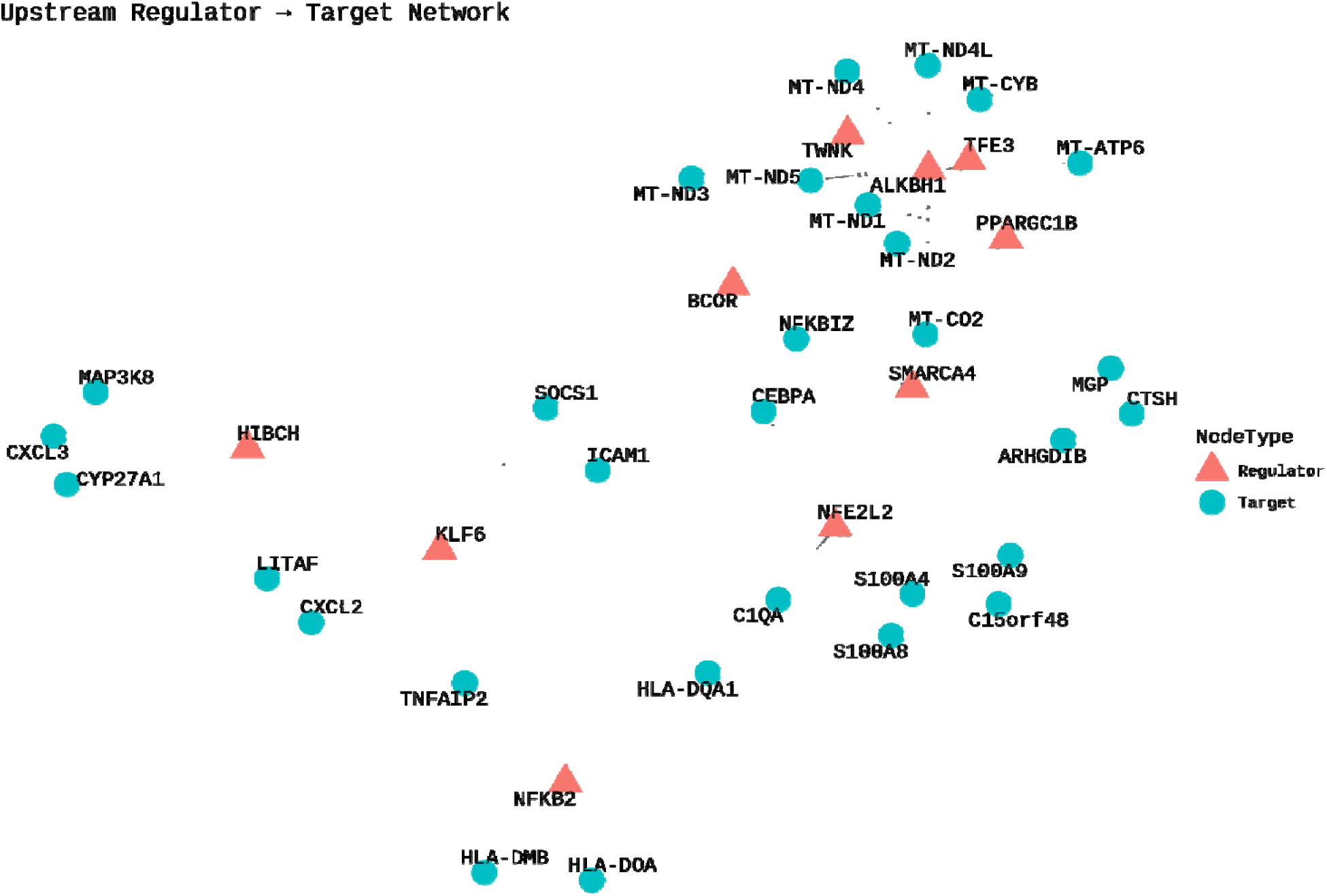

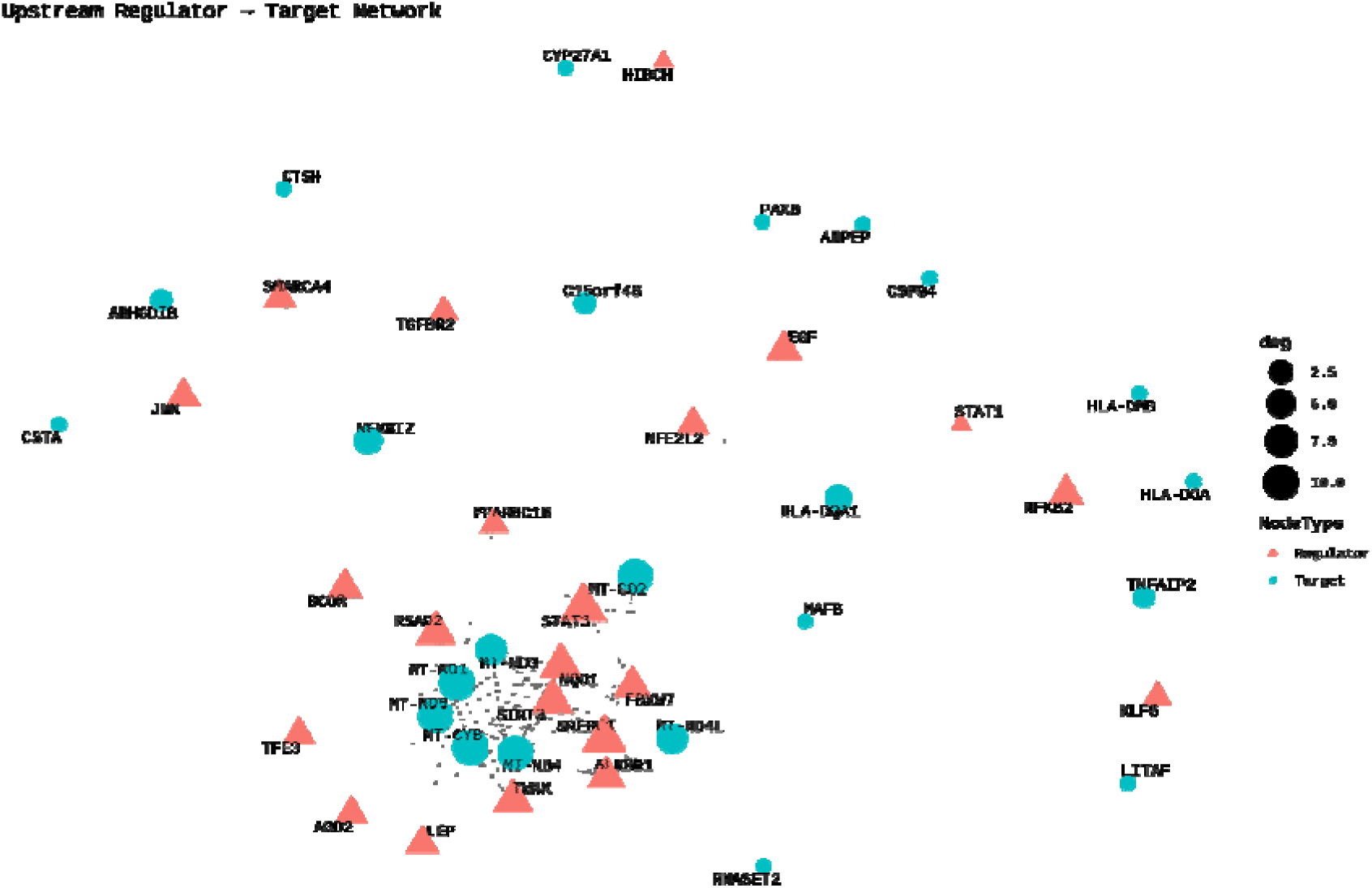

Additionally, we also looked into e novel regulatory interactions among downstream targets (Figure 4). The interactions revealed a coordinated **immunometabolic program** marked by mitochondrial remodeling and inflammatory activation. Several regulators were targeting **mitochondrial-encoded oxidative phosphorylation genes such as** (MT-ND1, MT-ND3, MT-ND4, MT-ND4L, MT-ND5, MT-CYB, MT-CO2). This regulation points to dysregulation in several processes such as mitochondrial respiration, redox balance, and mtDNA homeostasis. On the other hand the regulation by **NF-**κ**B, STAT, and TGF-**β**-associated regulators**, of antigen-presentation genes such as *NFKBIZ, TNFAIP2,* and *HLA-DQA1/DMB/DOA*, reflects a state of increased inflammation and immune-responsive.

**TABLE 2 of Upstream regulators and their novel interactions.**

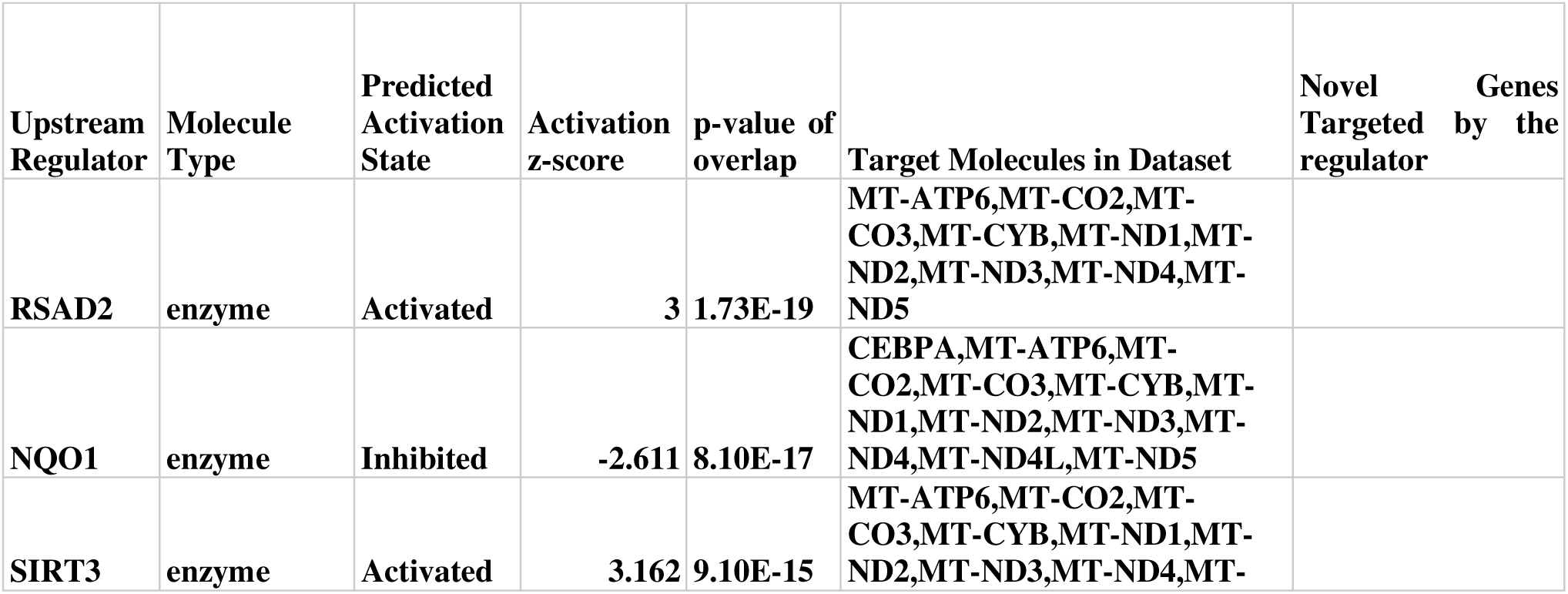
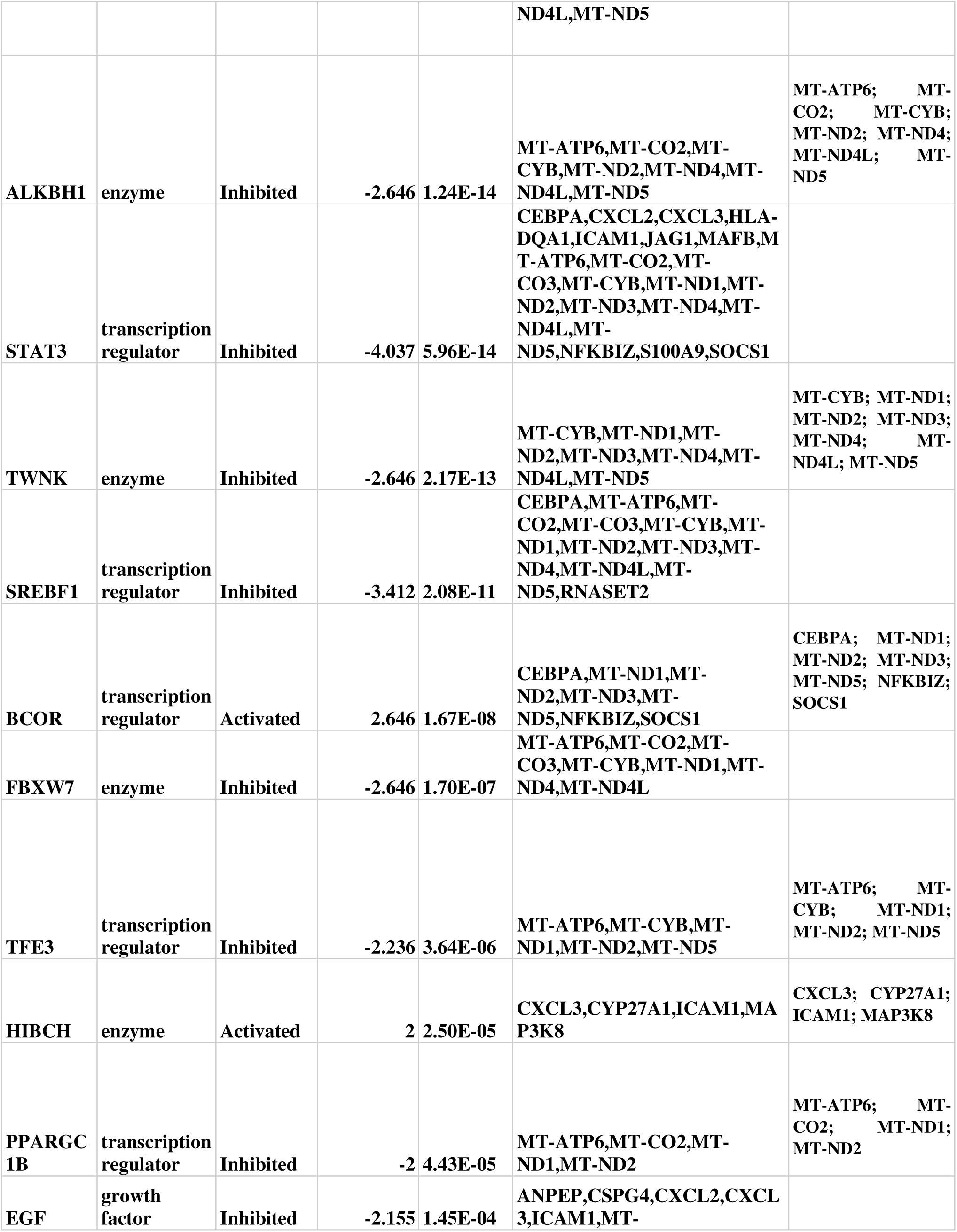
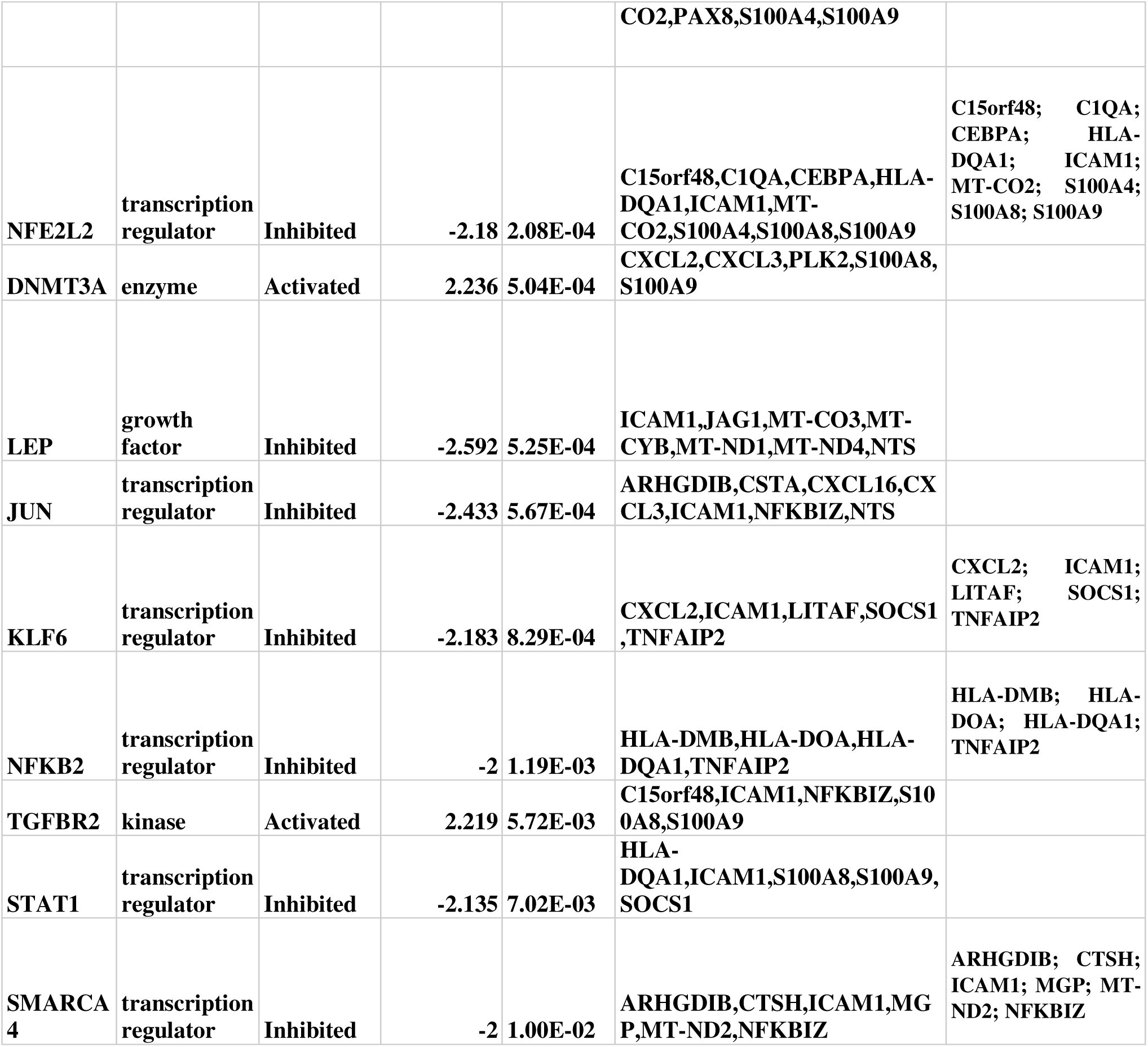

### Canonical Pathway Analysis

Ingenuity Pathway Analysis (IPA) revealed several canonical pathways as shown in **Figure 6 A and 6B**. Among the activated canonical pathways, were Granzyme A Signaling (p = 8E-02, z = 2.449) which showed upregulation of mitochondrial genes MT-ND1, MT-ND2, MT-ND3, MT-ND4, MT-ND4L, and MT-ND5, indicating increased mitochondrial activity associated with immune effector functions. Similarly, Parkinson’s Signaling Pathway (p = 1.9E-02, z = 2.449) and Sirtuin Signaling Pathway (p = 2.69E-02, z = 2.449) were activated with overlapping mitochondrial genes, suggesting upregulation of oxidative metabolism and stress response. The Mitochondrial Dysfunction pathway (p = 2.9E-02, z = 3.162) also emerged as significantly activated, involving MT-ATP6, MT-CO2, MT-CO3, MT-CYB, and multiple NADH dehydrogenase subunits (MT-ND1–ND5), consistent with increased mitochondrial bioenergetic activity and possible compensatory responses to stress or damage.

The inhibited pathways include the processes involving mitochondrial and immune dysfunction. tRNA processing in the mitochondrion (p = 2.44E-01, z = −3.162), rRNA processing (p = 3.12E-01, z = −3.162), and mitochondrial RNA degradation (p = 4.00E-01, z = −3.162), all shared subset of mitochondrial genes indicating coordinated downregulation of mitochondrial transcriptional and post-transcriptional machinery. Pathways related to oxidative phosphorylation (p = 9.26E-02, z = −3.162) and respiratory electron transport (p = 9.09E-02, z = −2.828) were also inhibited, suggesting a possible imbalance between mitochondrial biogenesis and energy production (38)(39). Inflammatory and immune pathways exhibited parallel suppression. Neutrophil degranulation (p = 1.89E-02, z = −3.0) and neutrophil extracellular trap signaling (p = 2.20E-02, z = −2.333) were notably downregulated, characterized by reduced expression of ADGRE5, ANPEP, CTSH, RNASET2, S100A8, and S100A9, reflecting diminished innate immune activation. Inhibition of hematoma resolution signaling (p = 3.09E-02, z = −2.828) and mitochondrial protein degradation (p = 5.10E-02, z = −2.236) further points to impaired tissue repair and mitochondrial quality control. Additional suppressed signaling cascades included Complex I biogenesis (p = 7.35E–02, z = −2.236), S100 family signaling (p = 6.39E-03, z = −2), Gα(i) signaling events (p = 1.55E-02, z = −2), RHO GTPase cycle (p = 8.89E-03, z = −2), and Class A/1 rhodopsin-like receptor signaling (p = 1.20E-02, z = −2). Antigen presentation pathways, including MHC class II and Class I MHC-mediated antigen processing, were also repressed (z = −2). Together these inhibited pathways indicate strong suppression of mitochondrial energy metabolism, cytoskeletal remodeling, G-protein–coupled receptor signaling, and innate immune activity, highlighting the shift toward metabolic adaptation and immune evasion.

Table 2 The novel canonical pathways associated with PCOS highlighted with *.

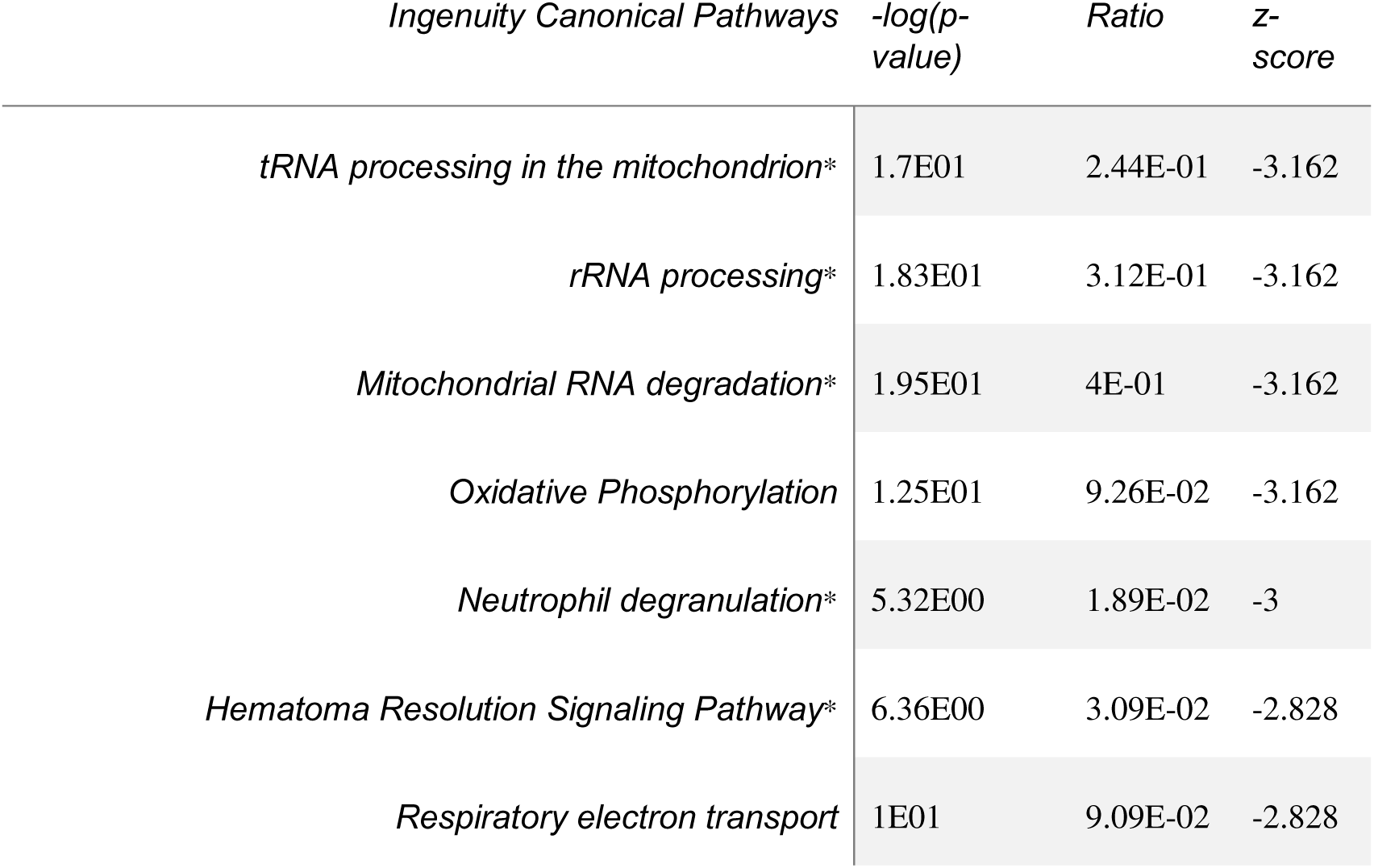
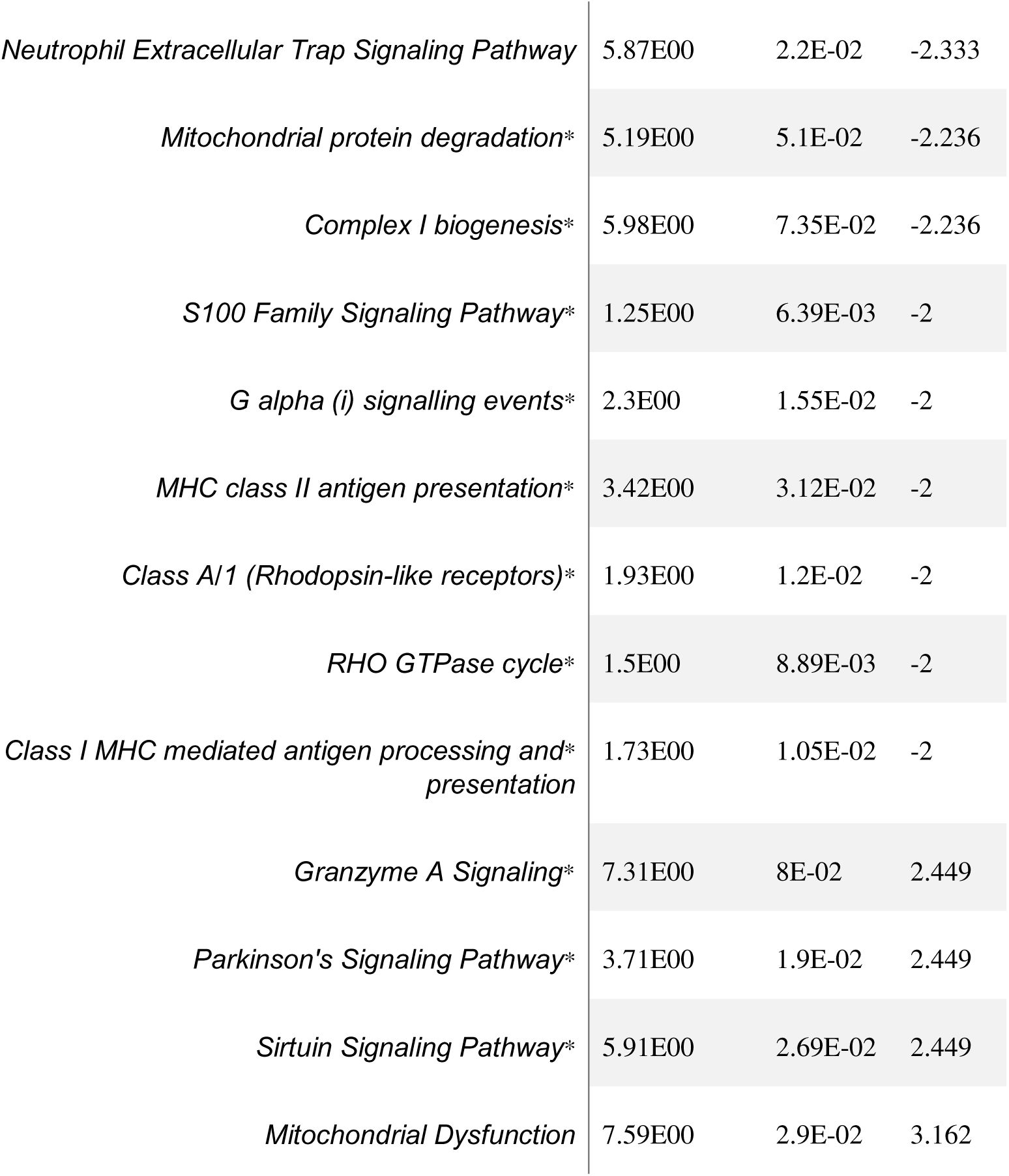

The new insights of of canonical pathway analysis characterizes PCOS **emerging issues of disruptions in mitochondrial quality-control pathways**, including mitochondrial protein degradation, RNA processing, and Complex I biogenesis table 2, rather than strictly associated with mitochondrial dysfunction and oxidative phosphorylation. These new insights into pathways suggests that impaired **mitochondrial maintenance and regeneration**, might provide a fundamental and underexplored mechanism in PCOS pathophysiology. Also, the increased immune and GPCR-related signaling pathways with limited association with PCOS suggests the impact of **immune-metabolic crosstalk and altered cellular signaling** in disease progression.

### Disease and Bio Functions

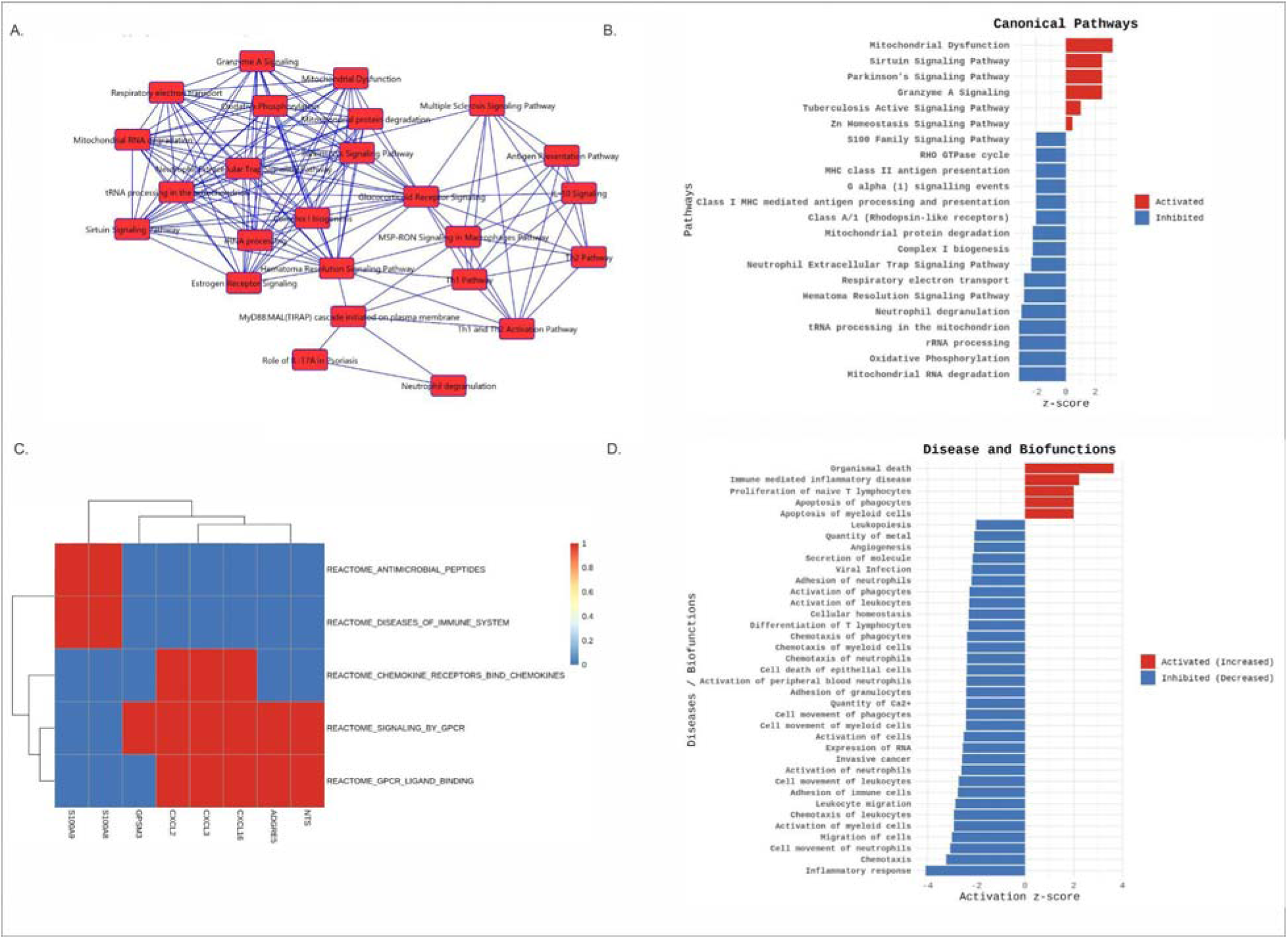

IPA analysis identified various biological functions, some activated and some inhibited Figure 6 D. The activated biofunctions were Proliferation of naïve T lymphocytes (z = +2.0), Apoptosis of phagocytes and myeloid cells (z = +2.0), Immune-mediated inflammatory illness (z = +2.219), and Organismal death (z = +3.642). These collectively signify elevated adaptive immunological activity, enhanced immune cell turnover, and stress-induced apoptosis. The association of ADGRE5, C15orf48, and SQOR in immune-mediated inflammation, and the role of ARHGAP45 and TAMALIN in T-cell proliferation and vesicular signaling, highlights the interaction between immune activation and mitochondrial redox adaptation. Furthermore, the activation of mitochondrial genes like ND4L and SQOR highlights a metabolic stress response that influences immunological regulation and apoptotic signaling (41). On the other hand, the inhibited biological functions suggest weakening of innate immune mechanisms, including inflammatory response (z = −4.087), chemotaxis and cell migration (z ≈ −3.24 to −3.01), activation of myeloid and neutrophil cells (z ≈ −2.9), and leukopoiesis (z = −2.02). The involvement of genes such as ADGRE5, ARHGAP45, ND4L, BOLA2B, and C15orf48 in these defined Biofunctions indicates compromised immune cell motility, adhesion, and differentiation, probably linked to mitochondrial dysfunction and altered redox equilibrium. The inhibition of cellular homeostasis (z = −2.33) and angiogenesis (z = −2.09), regulated by BOLA2B, SQOR, and TAMALIN, indicates a disrupted metabolic equilibrium and compromised abilities of for repair of the tissues (42) (43).The results showed that the processes of innate immune activation and immune cell trafficking were more prominent in PCOS patients, especially for neutrophils, myeloid cells, and phagocytes. The synchronized activation, chemotaxis, adhesion, and migration of these immune cell populations indicate that PCOS is linked to increased immune control and inflammatory cell accumulation, overcoming classical endocrine dysfunction. Several of these functions particularly neutrophil activation, extracellular migration, and phagocyte apoptosis present only little prior association with PCOS, highlighting them as novel immunological characteristics of PCOS. The simultaneous activation of leukopoiesis and T-cell differentiation indicates a more extensive dysregulation of hematopoietic and adaptive immune mechanisms, suggesting persistent immune remodeling rather than temporary inflammation. The involvement of calcium and metal ion transport, along with altered RNA expression, suggests potential connections between immune activation, intracellular signaling, and transcriptional regulation in PCOS.

## PPI NETWORK

The Protein-Protein Interaction (PPI) network consisted of 70 nodes and 61 edges, with the PPI enrichment p-value of (< 1.0 × 10□¹□). The protein interaction network is dominated by mitochondrial electron transport chain components, with MT-CO2 emerging as a central hub, indicating coordinated mitochondrial respiratory dysfunction figure 2C and 2D. The integration of PTGS1 within this network suggests a functional link between chronic inflammatory prostaglandin signaling and impaired mitochondrial bioenergetics in PCOS. Although mitochondrial dysfunction and inflammation are established features of PCOS, our network analysis provides novel integrative evidence linking PTGS1-mediated prostaglandin signaling with coordinated mitochondrial electron transport chain dysregulation, highlighting a previously underexplored inflammation-mitochondria axis in PCOS.

## GSEA Analysis

Gene Set Enrichment Analysis (GSEA) highlighted significant dysregulation of signaling and immune-related pathways in the polycystic ovary syndrome (PCOS) samples figure 7. Among the downregulated pathways were multiple G-protein-coupled receptor (GPCR)- associated pathways, including REACTOME_SIGNALING_BY_GPCR and REACTOME_GPCR_LIGAND_BINDING figure 7A and 7C. These enrichments indicate a significant suppression of receptor-mediated signaling involved in hormonal communication, neurotransmission, and metabolic regulation.

Immune-related pathways, such as chemokine signaling, antimicrobial responses, and immune system disorders, were also downregulated, suggesting impaired chemokine-mediated immune recruitment and weakened innate immune activation figure 7B and 7D. The genes contributing to these pathways include *CXCL2*, *CXCL3*, *CXCL16*, *ADGRE5*, *GPSM3*, *NTS*, *S100A8*, and *S100A9*. These genes are known as key regulators of immune cell trafficking, inflammatory signaling, and neutrophil function, supporting coordinated suppression of immune responsiveness in PCOS endometrium.

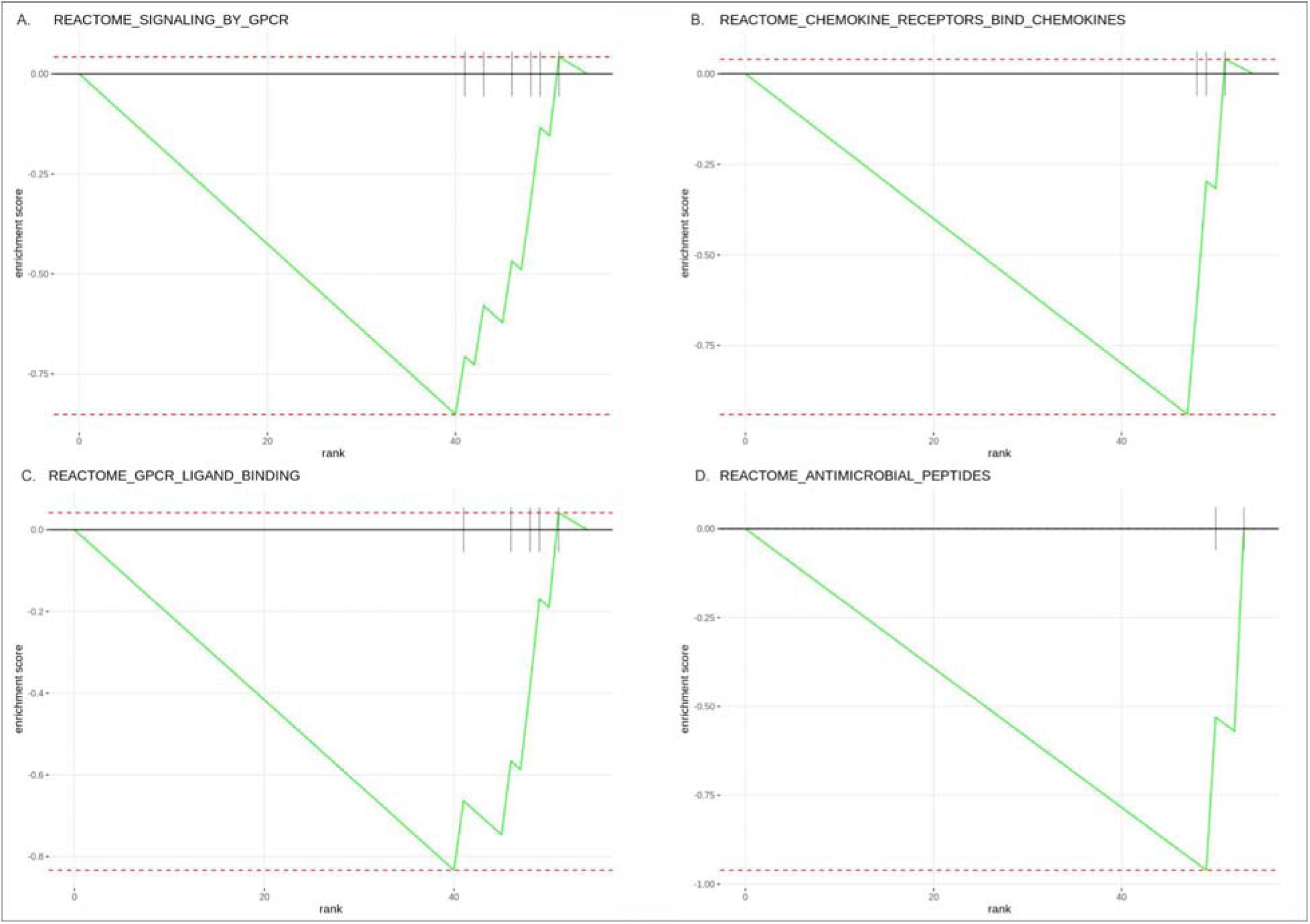

Several activated pathways were also highlighted which included the ones involved in glycosaminoglycan and chondroitin sulfate biosynthesis and metabolism. These enrichments suggest that there may be enhanced extracellular matrix biosynthetic activity and there is a shift toward structural remodeling suggesting changes in tissue architecture in response to altered immune and signaling homeostasis.

## Discussion

This study provides a comprehensive molecular interpretation of Polycystic Ovary Syndrome (PCOS), through novel insights involving genes such as C15orf48, ODF3B, PRR15-DT, LINC01176, and LOC105379193 that extend beyond the classical view of the disorder as an endocrine and insulin-resistance condition. In addition to the well-established hormonal and metabolic irregularities, our transcriptomic and pathway-level analyses highlight the persistent presence of chronic inflammation and oxidative stress (47) (48). The mitochondrial-immune convergence, characterized by altered expression of genes such as ND4L, SQOR, NFE2L2, C15orf48, and TGFBR2, which collectively link mitochondrial redox imbalance with innate immune modulation and fibrotic signaling suggests the role of mitochondrial dysfunction in PCOS (49). A thorough understanding of mitochondrial biology in PCOS reveals the role of mitochondrial dysfunction and oxidative stress as key features of the syndrome. Patients with PCOS consistently exhibit elevated mitochondrial reactive oxygen species (ROS) levels and disrupted bioenergetic levels (25). Insulin resistance and hyperandrogenism contribute to enhanced production of ROS, whereas antioxidant defense mechanisms appear depleted (47) (49). In the analysis, the dysregulation of mitochondrial genes such as ND4L and SQOR indicates impairment in electron transport and sulfide metabolism, validating established reports that connect PCOS to defective oxidative phosphorylation (OXPHOS) and increased oxidative stress. These molecular dysregulations suggest that PCOS cells involve some compensatory mechanisms, such as activation of sirtuin and antioxidant response pathways, to have minimal damage induced by bioenergetic failure.

The suppression of GPCR and chemokine signaling alongwith with selective adaptive activation, suggest reduced innate immune responses but elevated adaptive T-cell activity (50). The structural and biochemical reorganization is highlighted through extracellular matrix (ECM) and stromal remodeling. Further which is supported by disruptions in glycosaminoglycan and collagen turnover (51). Collectively, these observations support PCOS as an immunometabolic-mitochondrial disorder rather than a strictly endocrine condition. Moreover the observations also highlighted an interconnected mitochondrial-immune-fibrosis axis regulated by the interaction of NFE2L2 (NRF2), C15orf48 (MOCCI), and TGFBR2.

NFE2L2 serves as a master regulator of antioxidant defenses and becomes activated in response to oxidative stress, while C15orf48 encodes a mitochondrial open-reading frame that modulates inflammatory signaling (30). And TGFBR2 mediates TGF-β signaling, involved in fibrosis and stromal remodeling, which is frequently elevated in PCOS (52). Collectively, these molecules form an interconnected axis linking vital processes such as redox defense, immune response modulation, and fibrotic activity. The association of NRF2 and C15orf48, alongside the overactive TGF-β pathway, suggests that chronic oxidative stress in PCOS initiates adaptive mitochondrial responses and immunoregulatory adjustments, causing fibrotic remodeling within the tissue. These interactions align with prior evidence indicating that PCOS is marked by excessive TGF-β signaling, abnormal extracellular matrix deposition, and activation of oxidative stress-related transcription factors. Our findings also suggest that PCOS involves complex immune dysregulation and altered chemokine signaling, along with chronic low-grade inflammation and an imbalanced cytokine environment. The findings of gene ontology highlights **that there is a Immunometabolic reprogramming of the PCOS endometrium which is** characterized by **chronic cytokine-driven immune activation**,in addition to **nuclear/transcriptional remodeling and altered membrane signaling**. These biological functions are supported by **molecular functions linking oxidative stress sensing (RAGE), ion/metal regulation (calcium, zinc), and lipid-binding pathways**, collectively indicating that dysfunctionality in PCOS endometrial tissue arises from **integrated immune-metabolic stress rather than isolated endocrine disruption**.

Through IPA graphical summary a highly interconnected immune and inflammatory signaling network centered on **STAT3 was highlighted** showing that the gene STAT3 integrates signals from IL6, IL1B, TNF, IFNG, and IL17A suggesting the role of STAT3 as master regulator connecting cytokine signalling to PCOS. The involvement of Toll-like receptors (**TLR4, TLR7**) and MYD88 points to the activation of innate immune signalling pathways, which may sustain chronic inflammation and immune cell recruitment.

Upstream regulator analysis validated the involvement of regulators such as **SIRT3, STAT3, STAT1, SREBF1, and LEP with established association to** known roles of oxidative stress, lipid metabolism, and cytokine signaling. The identification of novel upstream regulators such as (**NFE2L2, TWNK, ALKBH1, BCOR, SMARCA4**) introduces some new insight into regulation of several processes including mitochondrial genome, redox balance, and chromatin remodeling. Canonical pathway analysis highlighted few activated and inhibited pathways. The activated pathways include the one related to mitochondrial dysfunction, sirtuin signaling, and stress responses, and whereas inhibited pathways involve the one essential in mitochondrial maintenance processes involving RNA processing, protein degradation, and Complex I biogenesis. This regulatory imbalance suggests that PCOS is not only characterized by altered energy metabolism but also by defective mitochondrial quality control and turnover.

The disease and Biofunction analysis highlighted activation of adaptive immune processes through T-cell proliferation and immune-mediated inflammation. The inhibited pathways included innate immune functions such as chemotaxis, myeloid activation, and leukopoiesis. These enrichments suggest that PCOS involves selective immune remodeling and favors chronic low-grade inflammation while dysregulating effective immune cell migration and tissue repair.

The PPI network analysis revealed an interaction which was mitochondrial-centered and dominated by electron transport chain components, with **MT-CO2** as a central hub. The integration of **PTGS1** with mitochondrial genes in the network provides novel evidence linking prostaglandin-mediated inflammation with mitochondrial respiratory dysfunction. This inflammation–mitochondria axis suggests that inflammatory signaling may directly impair bioenergetic capacity, strengthening metabolic stress in PCOS. Gene set enrichment analysis validated suppression of GPCR-mediated signaling and chemokine pathways. The suppression in the following emphasizes on the impaired hormonal communication and immune cell recruitment in PCOS endometrium. Also the extracellular matrix biosynthesis pathways points to the compensatory structural remodeling, suggesting adaptive changes to chronic inflammation and metabolic stress. Collectively, these observations state that PCOS a complex conditions rather than an endocrine condition characterized by **immunometabolic stress**, mitochondrial dysfunction, defective mitochondrial quality control, chronic yet dysregulated immune activation, epigenetic reprogramming, and altered tissue remodeling. This study has several limitations. It is based on a single publicly available endometrial dataset, so the results may reflect cohort-specific variation rather than universal features of PCOS. While multiple bioinformatic analyses were applied, the findings are observational and do not demonstrate causality. In addition, key clinical factors and technical influences, particularly those affecting mitochondrial gene expression, were not accounted for, and the proposed novel gene associations require validation in independent cohorts and experimental studies. Nonetheless, the integrative approach offers a useful framework to guide future mechanistic and validation research in PCOS.

## Conclusion

The findings of this transcriptomics study provides a comprehensive understanding that recharacterises Polycystic Ovary Syndrome (PCOS) as a complex condition. The condition marked by the immunometabolic-mitochondrial axis rather than being purely an endocrine condition. The DEG and pathway level analysis highlight that PCOS is characterized by coordinated mitochondrial dysfunction, oxidative stress, immune dysregulation, and tissue remodeling. The overlap of mitochondrial, immune, and fibrotic signaling through key regulators such as NFE2L2, C15orf48, TGFBR2, and STAT3, highlights a strong interconnected axis responsible for chronic inflammation, defective mitochondrial quality control, and extracellular matrix remodeling in the PCOS endometrium. The state of chronic inflammation with compromised immune recovery is highlighted through suppression of innate immune pathways alongside selective adaptive immune activation. Despite having limitations due to a single dataset, this study provides innovative insights and identifies potential regulatory targets that require validation in larger cohorts and experimental models, thus improving our understanding of PCOS pathophysiology.

## Acknowledgement

### Ethical Statement

This study does not contain any studies with human or animal subjects performed by any of the authors.

### Data Availability Statement

The data that support the findings of this study are openly available in [GEO] at <u>GEO Accession viewer</u>

### Author Contribution Statement

**Ritika:** Performing the complete end-to-end data analysis, writing the manuscript.

**Sonalika:** result interpretation

Prof R. C **Sobti:** reviewing the manuscript, providing critical feedback.

**Dr. Kashimir:** reviewing the manuscript, providing critical feedback.

